# A Scalable High Throughput Fully Automated Pipeline for the Quantification of Amyloid Pathology in Alzheimer’s Disease using Deep Learning Algorithms

**DOI:** 10.1101/2023.05.19.541376

**Authors:** Vivek Gopal Ramaswamy, Monika Ahirwar, Genadi Ryan, Maxim Signaevsky, Vahram Haroutunian, Steven Finkbeiner

**Affiliations:** Center for Systems and Therapeutics and Taube/Koret Center for Neurodegenerative Research, J. David Gladstone Institutes, San Francisco, California 94158; Department of Pathology, Icahn School of Medicine at Mount Sinai, 1 Gustave L. Levy Place, Box 1194, New York, NY, 10029, USA; Nash Family Department of Neuroscience, Icahn School of Medicine at Mount Sinai, New York, New York, USA; Department of Psychiatry, Icahn School of Medicine at Mount Sinai, New York, New York, USA.; Alzheimer’s Disease Research Center, Icahn School of Medicine at Mount Sinai, New York, New York; JJ Peters VA Medical Center - MIRECC, Bronx, New York, USA; Departments of Neurology and Physiology, University of California, San Francisco, San Francisco, California 94158

## Abstract

The most common approach to characterize neuropathology in Alzheimer’s disease (AD) involves a manual survey and inspection by an expert neuropathologist of postmortem tissue that has been immunolabeled to visualize the presence of amyloid β in plaques and around blood vessels and neurofibrillary tangles of the tau protein. In the case of amyloid β pathology, a semiquantitative score is given that is based on areas of densest pathology. The approach has been well-validated but the process is laborious and time consuming, and inherently susceptible to intra- and inter-observer variability. Moreover, the tremendous growth in genetic, transcriptomic and proteomic data from AD patients has created new opportunities to link clinical features of AD to molecular pathogenesis through pathology, but the lack of high throughput quantitative and comprehensive approaches to assess neuropathology limits the associations that can be discovered. To address these limitations, we designed a computational pipeline to analyze postmortem tissue from AD patients in a fully automated, unbiased and high throughput manner. We used deep learning to train algorithms with a Mask Regional-Convolutional Neural Network to detect and classify different types of amyloid pathology with human level accuracy. After training on pathology slides from a Mt Sinai cohort, our algorithms identified amyloid pathology in samples made at an independent brain bank and from an unrelated cohort of patients, indicating that the algorithms were detecting reproducible and generalizable pathology features. We designed the pipeline to retain the position of the pathology it detects, making it possible to reconstruct a map of pathology across the entire whole slide image, facilitating neuropathological analyses at multiple scales. Quantitative measurements of amyloid pathology correlated positively and significantly with the severity of AD as measured by standard approaches. We conclude that we have developed a computational pipeline to analyze digitized images of neuropathology in high throughput and algorithms to detect types of amyloid pathology with human level accuracy that should enable neuropathological analysis of large tissue collections and integration of those results with orthogonal clinical and multiomic measurements.

## Introduction

Alzheimer disease (AD) is the most common adult onset neurodegenerative disease. It is characterized by a devastating loss of memory and executive function and is frequently associated with behavioral changes^1, 2^. Human genetics studies have provided critical insights into the molecular underpinnings of AD^3^. Rare damaging and protective mutations in the gene encoding the amyloid precursor protein (*APP*) and in enzymes responsible for its proteolysis and metabolism have established that it plays a causative role in AD^4–14^. Alleles of apolipoprotein E (*APOE*), a lipid binding protein, are the greatest genetic risk factor for AD, but rare variants in the transmembrane receptor expressed in cells of the myeloid lineage (*TREM2*) also have large effects on AD risk^15^. In recent work, 75 additional genetic loci have been associated with AD risk at a genome-wide level^3^. The initial analysis of these loci strongly implicates microglial dysfunction, neuroinflammation, and the autophagy/endolysosomal signaling pathway, in addition to amyloid/tau pathways, as contributors to AD. Overall, while the major symptoms of AD can be explained by the dysfunction and loss of neurons, the genetics strongly suggest that AD is a complex emergent property involving interactions between neurons and many other cell types.

Neuropathological assessment of brain tissue from AD patients confirms numerous abnormalities that correlate with genetic findings. The abnormal deposition of amyloid β, a cleavage product of the APP protein, and the presence of neurofibrillary tangles (NFTs) of hyperphosphorylated tau, are the two diagnostic histopathological hallmarks of AD. But other prominent abnormalities include the preferential loss of neurons from different brain regions, microglial activation, astrocytosis, vascular and white matter changes, and the deposition of other proteins including a-synuclein and Tar DNA binding protein 43 (TDP-43)^16–18^. It may be necessary to appraise all these features at once to understand whether and how they contribute to AD symptom profile and progression and to identify reliable markers or marker combinations that predict disease risk, type and stage, since canonical markers of AD such as amyloid β plaques and NFTs, as currently used, appear to only have limited predictive value.

The first descriptions of plaques in brain tissue from people with AD were reported in the early 20^th^ century, and amyloid β was established as a component of plaques in the mid 1980s. But the role of amyloid β and plaques in AD has been debated since their discovery. A dominant idea in the field, encapsulated by the amyloid hypothesis, posited that these plaques were a driving factor in AD pathogenesis^19^. The fact that these structures appeared early in the course of AD and often were closely associated with activated microglia added circumstantial evidence for their role^20^. On the other hand, amyloid β plaques are observed in cognitively normal individuals, and the amount of amyloid β pathology correlates relatively poorly with symptom profile. The role of amyloid β plaques was cast in further doubt when clinical trials found that several agents that substantially increased their clearance in patient brains did not lead to clinical improvement, and even worsened the symptoms of some patients^21, 22^. On the other hand, two antibody therapies directed against amyloid β were recently shown to induce modest but statistically significant improvements in clinical endpoints in AD patients in addition to reducing amyloid plaque pathology in patient brains^23^. These contradictory findings underscore the importance of better defining the nature of amyloid β pathology that drives AD. Our current hypothesis is that amyloid pathology is a defining determinant of AD, but that finding reliable correlations between pathology and disease requires methods that are more quantitative, comprehensive, and sensitive than current rating systems, and that can detect more specific features of amyloid pathology than currently possible.

Conventionally, amyloid pathology in patient brain tissue has been assessed with immunolabeling and visual microscopic inspection by expert neuropathologists. In 1991, the Consortium to Establish a Registry for Alzheimer Disease (CERAD) proposed diagnostic criteria for AD that used a semi-quantitative score of the density of neuritic plaques in the most severely affected regions of the isocortex, and the patient’s age at death to obtain an age-adjusted plaque score^24^. This semi-quantitative approach to plaque quantification continues, even as the specificity of the neuropathological diagnosis of AD has been improved by incorporating Braak and Braak staging of NFTs. More recently, new approaches to quantify amyloid β plaques are being developed that utilize machine learning and computer vision. Several groups have shown on relatively small samples that it is possible to train deep learning algorithms to achieve human level accuracy identifying amyloid β pathology^25–28^.

The ability to quantify amyloid β pathology accurately, rapidly, inexpensively, comprehensively and in an unbiased and automated manner would be a major achievement. It might make it possible to scale neuropathological analyses and generate datasets of sufficient size to do meaningful association studies with multiomic data from patients. Such studies could in turn lead to the discovery of new causal genetic variants, elucidate genotype-phenotype relationships, gauge therapeutic responses, and reveal the nature of the heterogeneity in clinical profiles of AD patients. The studies utilizing machine learning have served as proof-of-concept for the approach and even demonstrated improvements over the conventional CERAD scoring system^29^.

However, many challenges remain. First, digitization of pathology slides creates enormous electronic files that are difficult to store and process with machine learning pipelines. These whole slide images (WSI) have a lot of variability in them induced by stain intensity, tissue preparation and slide scanning techniques. To ingest the imaging data, WSIs are often divided into smaller crops that are easier to compute. But the detection and interpretation of neuropathology often depends critically on context. If the crops are too small, some pathology could be overlooked or misdiagnosed. Second, the success of supervised machine learning approaches depends critically on the production of well-annotated training datasets. Training datasets provide the ground truth from which the machine learning network can learn. They are typically generated by expert neuropathologists who hand-annotate examples of characteristic pathology vs. unspecific staining in digitized images. But hand annotation is time consuming and expensive, and so it is often difficult to generate sufficiently large training datasets to produce accurate algorithms. Lastly, many processing pipelines are trained to recognize and count specific types of neuropathology without segmenting the pathology into individual pixels. The challenge posed by amyloid β plaque pathology illustrates why this is an important limitation: simple counts of plaques do not correlate well with the severity of AD dementia, yet antibodies targeting amyloid β appear to improve clinical endpoints. Tools that segment amyloid β pathology might uncover subtle features in the plaques, such as their shape, size, texture, tissue context, etc…, that more directly relate to AD than plaque number.

To address these limitations, we designed a computational pipeline that can process digital neuropathology images in high throughput, and is scalable to large datasets. Importantly, we designed it so that it not only identifies neuropathology but simultaneously segments neuropathological objects for deeper types of analyses. We then trained this pipeline to recognize key types of amyloid β pathology, including cored and diffuse amyloid β plaques as well as cerebral amyloid angiopathy, with human level accuracy. The models performed well using standard tests, and critically, generalized to unseen data from an independent data set. In addition, we demonstrated a positive correlation between various measures of amyloid β pathology using these automated analyses and the CERAD score of AD severity for each patient.

## Results

### Improving throughput, accuracy and depth of analysis

We designed our machine learning computational pipeline and workflow for the analysis of digitized pathology samples with several goals in mind (Fig. 1). First, we wanted to improve throughput to make it feasible to perform analysis of large brain bank collections at scale and to better integrate the results with clinical and genomic data. Whole slide images (WSI) are very large files (100,000 x 100,000 pixels), and batch processing large sets of WSIs can cause premature job termination due to timeout errors. WSIs are also too large to ingest into deep learning (DL) networks, so they are typically divided into tiles 256 x 256 pixels in size, which takes approximately 109 minutes per WSI. To optimize throughput, we increased the crop size to 1024 x 1024 pixels (Fig. 1, Step 3). By reducing the number of tiles per WSIs by 16-fold, the pipeline is able to process an entire WSI in less than 21 minutes. Nevertheless, many pathological hallmarks are sparsely distributed, and so many crops lack positive features. To avoid wasting computational time analyzing crops that lack important features, we added a regional proposal network (RPN) to the architecture of the convolutional neural network, thus termed R-CNN. The RPN slides a small network window over the convolutional feature map put out by the last layer of the CNN, and determines the probability that the window contains an object (Fig.1, Step 4; Fig. 11, Step 2). The proposed regions with a high likelihood of containing objects of interest are then fully analyzed by the CNN (Fig. 1, Steps 5-6; Fig. 11 Step 2), whereas those with low likelihood are ignored, saving a tremendous amount of compute time.

**Figure 1.**
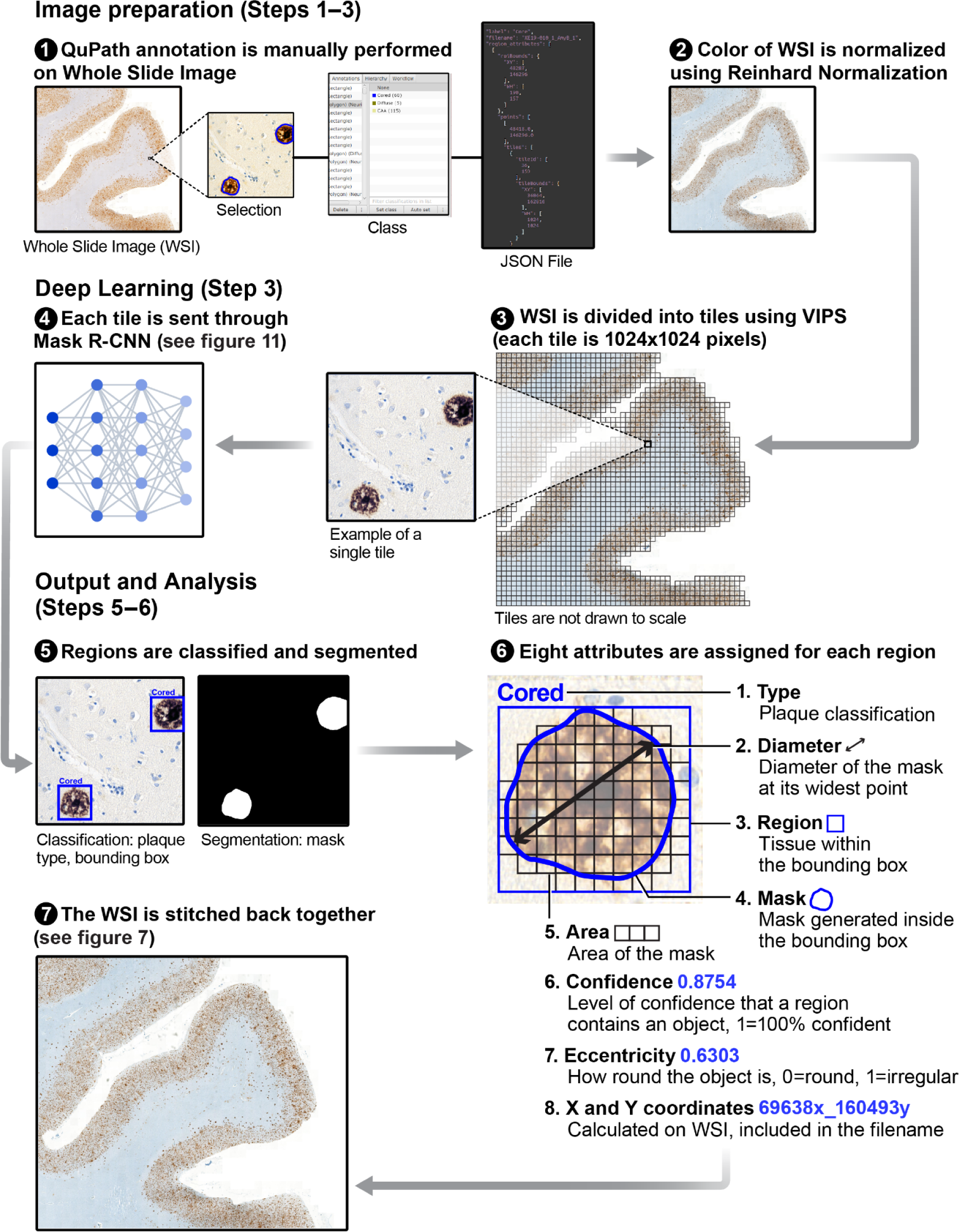
High-throughput and scalable pipeline for digital pathology analysis.

Second, we wanted to improve the accuracy of the pathology analyses. Context is one important factor that expert pathologists use to determine accurately the nature of microscopic pathology. So, another motivation for increasing the size of the tiles from 256 x 256 to 1024 x 1024 was to provide greater context to improve the accuracy of object classification.

Third, we wanted to increase the depth of the analysis. To this end, we implemented a Mask R-CNN^30^. Image analysis by Mask R-CNN goes beyond simple object identification and performs object detection and segmentation simultaneously by sharing CNN weights. The availability of automated object segmentation opens up an enormous array of options to analyze features within the pathological objects, such as amyloid plaques or neurofibrillary tangles, that are not possible otherwise, and may lead to the discovery of more specific features of these objects with particular diagnostic or prognostic significance.

To perform supervised DL with the Mask R-CNN, we first need to develop a training dataset that includes examples of pathology we want the Mask R-CNN to learn to recognize and examples of nonspecific labeling we want it to ignore. To develop those training datasets, we depend on expert neuropathologists to generate ground truth annotation of examples within the dataset. To make the annotation process easier, we developed a custom script that helps pathologists rapidly annotate regions of interest over large context areas in QuPath Software, an open-source software for digital pathology image analysis^31^.

### Precise localization and segmentation of Aβ IHC-positive histopathological diagnostic elements

Having constructed a computational pipeline for the analysis of WSIs, we used it to train a Mask R-CNN-based model to classify and segment **Aβ IHC-positive histopathological diagnostic objects. This category comprises Amyloid plaques and pathological Aβ accumulation in vessel walls constituting** cerebral amyloid angiopathy (CAA). For ground truth annotation purposes we developed a set of operational criteria based on widely accepted classifications and definitions used for diagnostic and machine learning purposes. For Amyloid plaques we based our operational morphological criteria in alignment with the recent studies done by Boon *et al*., 2020^32^ and DeTure *et al*., 2019^17^. Briefly, for ground truth annotation and further training purposes we defined *Diffuse* plaques as poorly marginated collections of Aβ IHC-positive lacy staining lacking dystrophic neurites and organized internal structure; *Cored* plaques - as the one with the central dense core surrounded by an empty corona or halo and further a ring of diffuse Aβ IHC-positive lacy staining with or without dystrophic neurites; *Coarse-grained* plaques - as these with ill-defined borders, with dystrophic neurites and small dense cores but lacking one central core and an empty halo; CAA, **which is an important well described and not an uncommon comorbidity of AD**^33^ is characterized by the accumulation of Aβ deposits within the walls of small to medium-sized cerebral blood vessels^34, 35^.

The training dataset consisted of 3188 tiles (1024×1024 pixels each) from 10 WSIs. We ensured that the training tiles did not always contain the objects of interest at their center, so the model would not erroneously learn to concentrate on the tiles’ central area. The tiles were annotated by an expert pathologist, who provided both a classification label and a pixel-level segmentation mask for each object in QuPath. After training, Mask R-CNN was evaluated on a validation dataset consisting of a different set of 7 WSIs. These WSIs had also been annotated by an expert. During the validation process, the model’s predictions on these WSIs are compared against the expert annotations to assess accuracy. The trained Mask R-CNN predicted both the classification labels and the corresponding segmentation masks for each individual tile (Fig. 2a) with human-level accuracy. In addition, Mask R-CNN was able to classify and segment multiple pathological objects within a single tile (Fig. 2b), which is important for achieving comprehensive and accurate quantification of neuropathology from the 1024×1024 crops. Finally, we note that amyloid β can be found in the image outside the objects of interest.

**Figure 2.**
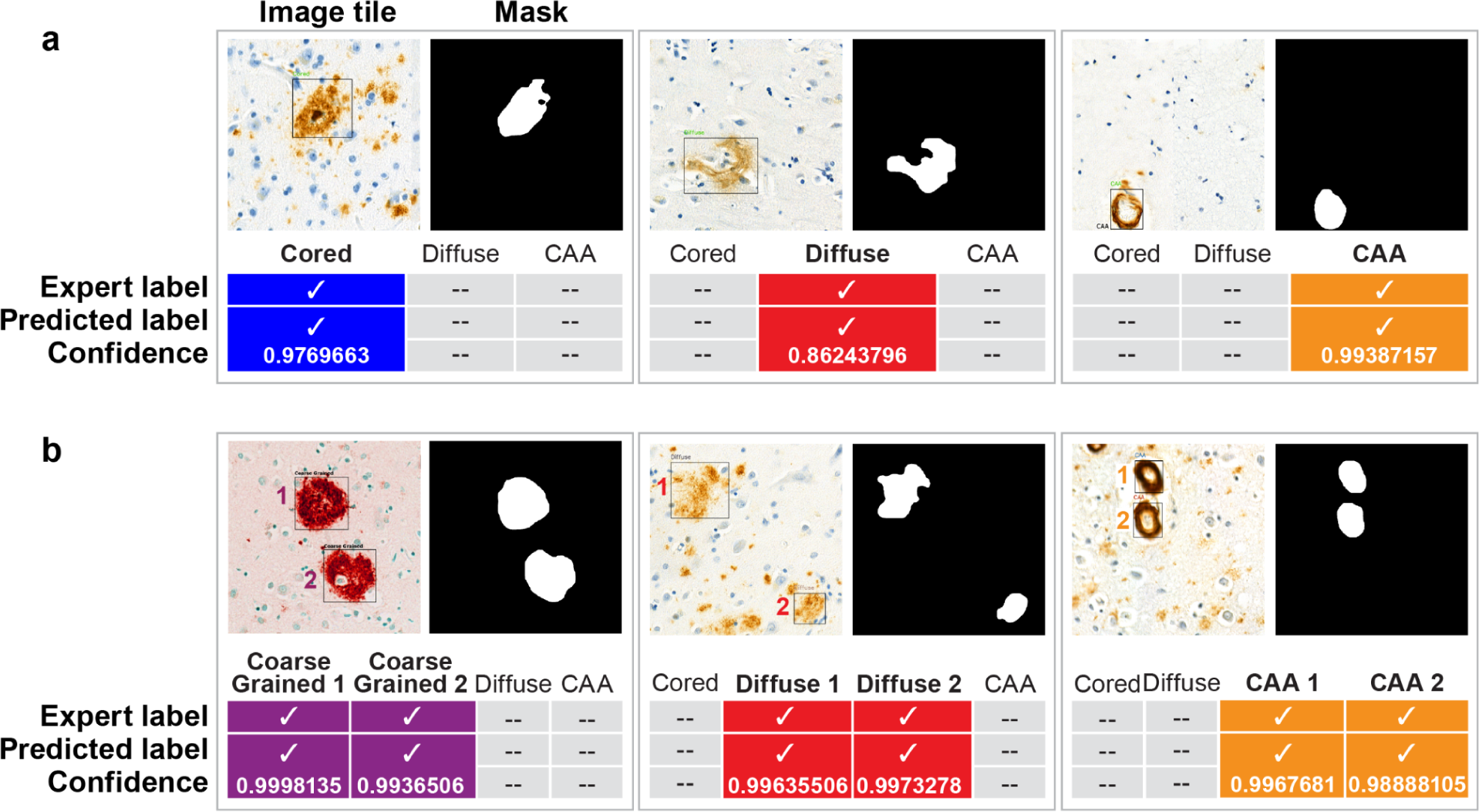
A Mask R-CNN was trained to accurately predict different types of amyloid neuropathology. a) Model output for the validation set obtained from the Mount Sinai NIH Brain and Tissue Repository. For each tile, both the expert label and the predicted label are shown. Each predicted label also has a confidence score where 1 corresponds to 100% confidence. b) Nevertheless, the model can detect, classify and segment multiple objects in a single tile.

Just as the DL network is trained by example to learn the amyloid β that forms pathology recognized by pathologists, the DL network also learns to ignore non-specific amyloid β labeling that does not conform to defined classes of pathology.

## Model Testing, Validation and Generalizability

### Mask R-CNN evaluated on a histology dataset surpasses the benchmark performance of Mask R-CNN evaluated on the COCO dataset

To evaluate our model’s performance, we employed the evaluation metrics provided by Microsoft’s Common Objects in Context (COCO)^36^. The COCO benchmark is considered a state-of-the-art technique for assessing the capabilities of object detection and segmentation models. It offers standardized techniques and a leaderboard for comparing and benchmarking performance across different models. The COCO benchmark has 12 metrics, calculating Average Precision and Average Recall at a series of IoU (Intersection over Union) thresholds and across varying object sizes (Table 1). Precision, recall and IoU are explained below.

**Table 1.** 12 benchmark metrics used for characterizing the performance of an object detector on COCO.

|  |  |
| --- | --- |
| Average Precision (AP) | AP = % of true positive predictions relative to all the predicted detections<br>AP : AP at IoU=.50:.05:.95 (**primary challenge metric**)<br>AP<sub>IoU=.50</sub> : AP at IoU=.50 (PASCAL VOC metric)<br>AP<sub>IoU=.75</sub> : AP at IoU=.75 (strict metric) |
| AP Across Scales | AP<sub>small</sub> : AP for small objects: area < 32<sup>2</sup><br>AP<sub>medium</sub> : AP for medium objects: 32<sup>2</sup> < area < 96<sup>2</sup><br>AP<sub>large</sub> : AP for large objects: area >96<sup>2</sup> |
| Average Recall (AR) | AR = % of true positive predictions relative to the ground truth objects<br>AR<sub>max=1</sub> : AR given 1 detection per image<br>AR<sub>max=10</sub> : AR given 10 detections per image<br>AR<sub>max=100</sub> : AR given 100 detections per image |
| AR Across Scales | AR<sub>small</sub> : AR for small objects: area < 32<sup>2</sup><br>AR<sub>medium</sub> : AR for medium objects: 32<sup>2</sup> << area < 96<sup>2</sup><br>AR<sub>large</sub> : AR for large objects: area >96<sup>2</sup> |

When performing object detection, the model draws a bounding box around each detected object in the predicted dataset (Fig. 1, Step 5). Similarly, the annotator draws a bounding box around each object in the ground-truth dataset. IoU is the overlap between the bounding box around the predicted object and the bounding box around the ground-truth object. It is calculated by dividing the area of intersection between the predicted and ground-truth boxes by the area of their union. A higher IoU indicates a better match and implies more accurate detection. Object detections with an IoU below a set threshold are rejected.

Precision measures how well the object detection system correctly identifies objects within these bounding boxes. It is determined as the proportion of true positive predictions (correctly identified objects) among all the predicted detections. Recall refers to the ability of a system to detect all instances of a particular object within an image. It is determined as the proportion of true positive detections with respect to the total number of ground truth objects at each IoU threshold.

Setting the IoU threshold at 0.5 (default value for Mask R-CNN) means that any predicted mask that overlaps with the ground-truth mask with an IoU of 0.5 or more is considered a true positive. A higher IoU threshold increases precision because it only accepts as truly positive the predicted masks that have a large overlap with the ground-truth masks. Many papers compute Average Precision at a specific IoU threshold (typically at 0.5) to evaluate their object detection system. Instead, we averaged the AP over IoU thresholds ranging from 0.5 to 0.95 in increments of 0.05. By calculating AP at multiple IoU thresholds (e.g., 0.5, 0.55, 0.6, etc.), we can detect strengths and weaknesses of the model at different levels of object overlap, incorporating the variation in performance metrics. IoU thresholding also controls the trade-off between precision and recall, thus using a range of IoU thresholds will help the model handle this trade-off and its overall performance at different precision-recall levels. Thus our evaluation metric provides a robust assessment of the model’s ability to accurately detect objects with different degrees of precision.

The model achieved an average precision of 0.422 for object detection, and an average precision of 0.401 for segmentation (Table 2). We compared this performance to that of a Mask R-CNN model evaluated on the COCO minval dataset consisting of 5000 validation images. Our model exceeded the benchmark performance of the Mask R-CNN model by 11.93% on the average precision for object detection and 15.90% for object segmentation.

**Table 2.** Model performance comparison with object detection and segmentation benchmark. The boxAP and maskAP are defined as AP @[IoU=0.50:0.95| area=All|maxDets=100].

| Model | boxAP | maskAP |
| --- | --- | --- |
| Mask R-CNN trained on COCO dataset | 37.7 | 34.6 |
| Mask R-CNN trained on AD histopathology WSI dataset | 42.2 | 40.1 |

As further evaluation of our model’s performance, we calculated the precision-recall curve for all object classes at 0.5 IoU threshold (Fig 3). An ideal model would have a precision and recall of 1 at all IoUs. A Precision-Recall curve is considered good when it lies on the top left side of the diagonal line. Our model showed a good balance between precision (percentage of true positives in total predicted objects) and recall (percentage of true positives in ground truth objects), as our precision stayed high as recall increased.

**Figure 3.**
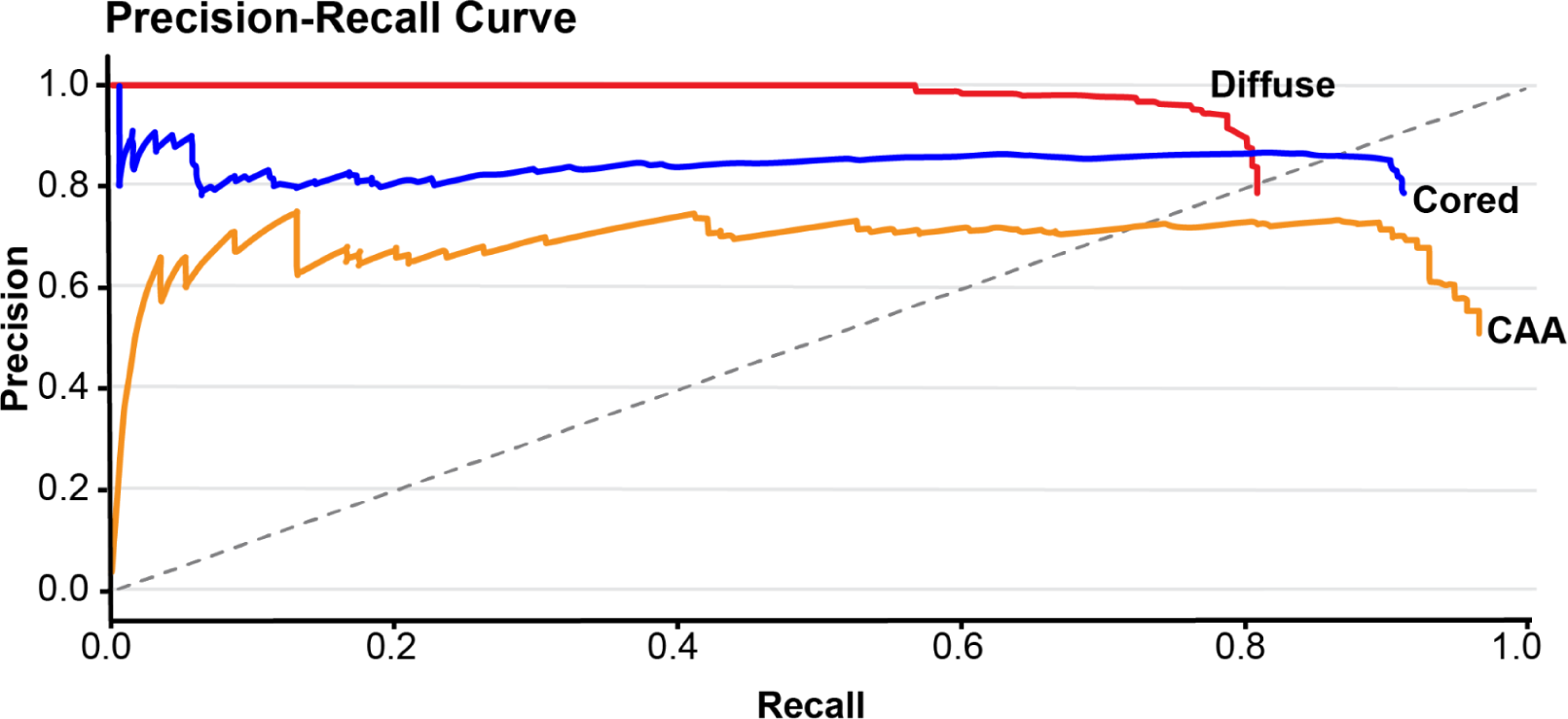
The Mask R-CNN models trained to predict different types of amyloid pathology performed well based on precision-recall metrics. a) Shows a Precision-Recall Curve for validation dataset from Mount Sinai NIH Brain and Tissue Repository for all the three classes - Cored, Diffuse and CAA - at 0.5 IoU threshold. A curve closer to the top-left corner indicates better model performance.

### The trained Mask R-CNN is not overfitted to the training dataset

One challenge with training deep learning models is that the technique is so sensitive that the models can become overfit, which means that the model learns to make classifications based at least partly on peculiarities in the training data set that do not generalize. Consequently, the accuracy of an overfit model falls when it is applied to unseen data, limiting its utility. A common method for assessing if a model is overfit is k-fold cross validation, which entails first dividing the dataset into k non-overlapping sets, and then training k models iteratively using the (k-1) sets as the training set and remaining one as the validation set. The aggregated accuracy of these k models provides a better estimate of model’s performance, which further helps in assessing how well the model generalizes to new data and reduces the risk of overfitting or underfitting. For our k-fold testing, we combined the 10 WSIs from the training dataset with the 7 WSIs from the validation dataset, and broke them down into k non-overlapping data subsets.

Choosing an appropriate value of k in k-fold cross-validation is important because it can impact the reliability of the performance estimates and the computational cost of the evaluation process. Selection of k depends upon the size of the dataset, available computational resources, and the bias-variance trade-off. We picked four values of k (3, 5, 7 and 9), and for each value, we asked the k models to detect and segment the three classes of beta-amyloid pathology. We used the AP averaged over IoU thresholds between 0.5 to 0.95 in increments of 0.05 to evaluate the performance of all the models at each k value, and repeated the process five times to calculate bias and variance. We observed that with increasing value of k, our bias and variance decreased (Fig 4). This is expected because with k = 3, we fit the models using 2/3 of the dataset for training, and 1/3 for validation giving higher variance and bias, but with k = 9, we fit each model on a much bigger dataset 8/9 for training, and 1/9 for validation giving less bias and variance. For k = 5, 7 and 9 we obtained comparable mean AP for both segmentation and boundary box detection (Fig. 4 a, c). In addition, the deviation from mean AP was not significant for either bounding box prediction or segmentation (Fig 4 b,d).

**Figure 4.** K-fold cross validation testing indicates model performance remains relatively stable across different data segments. K-Fold cross validation performed on the dataset from Mount Sinai NIH Brain and Tissue Repository. a) Box plot for AP @[IoU=0.50:095| area=All|maxDets=100] vs k, where value of k is {3, 5, 6, 9} b) Percentage change in (AP) @[IoU=0.50:095| area=All|maxDets=100] wrt. mean for each k {3, 5, 7, 9} for object detections. c) (AP) @[IoU=0.50:095| area=All|maxDets=100] vs k, boxplots obtained by simulating 5 runs for each k{3, 5, 7, 9} for segmentation. d) Percentage change in (AP) @[IoU=0.50:095| area=All|maxDets=100] wrt mean for each k {3, 5, 7, 9} for segmentation. At k=5, 7, 9 we obtain comparable mean AP for both segmentation and object detection.

If a model performs differently on different datasets, it indicates that the model has learned some irrelevant patterns or noise specific to its training set, and struggles to perform well when faced with new datasets. The fact that at k = 5 and above, the AP of our models only varied by +/- 0.2 or less indicates that they are not overfitted to their training set.

Given that variance and bias stabilized by k = 5, and given the cost of computation for increased values of k, we consider that k = 5 is a good value for k-fold cross evaluation of our type of model on our type of dataset. At this value of k, the average AP values were 0.54+/-0.2 for boundary box prediction, and 0.51+/-0.2 for segmentation. These values represent our model’s assessment for new data, and indicate a low risk of overfitting.

### Mask R-CNN generalizes well to an independent biological dataset

To assess the generalizability of the Mask R-CNN model beyond the dataset that was used to train it, we applied it to a completely independent dataset, AD neuropathology samples from a different cohort of patients and from an unrelated brain bank at the University of California, Davis. A total of 160K tiles were extracted from the UC Davis dataset (external). This external dataset differs from our dataset by its tile size (256 x 256 instead of 1024 x 1024) and reference normalization image, and by the fact that each tile is assigned a single class - Cored, Diffuse and Neuritic (whereas our larger tiles can have a mixed class label). Using our Mask R-CNN model on this external dataset, we obtained 71% accuracy on Cored, 57% accuracy on Diffuse and 76% accuracy on CAA at 0.5 IoU threshold. The Precision-Recall curves were somewhat better for Cored and CAA than Diffuse classes (Fig. 5a). The Receiver Operating Characteristic (ROC) Curve, a measure of a model’s ability to distinguish classes, indicated more accurate predictions for Cored than other classes (Fig 5 b). That our models performed well despite the fact that training and testing datasets differed in format (tile size), and biological sample preparation and labeling, indicates that the models have learned generalizable features of amyloid pathology.

**Figure 5.**
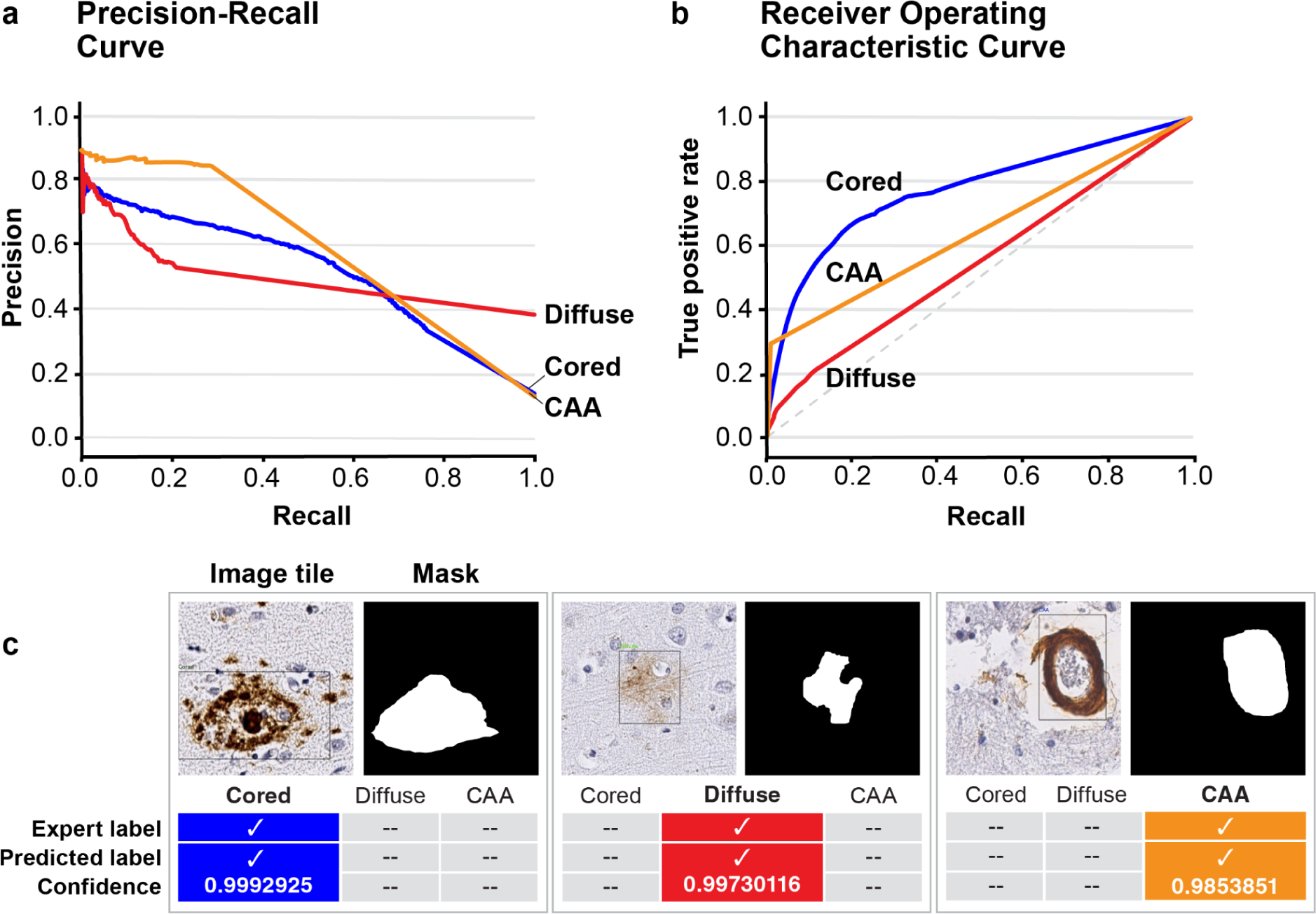
External data validation a) Precision Recall Curve for external dataset with for all the 3 classes - Cored (Blue line), Diffuse (Red line) and CAA (Green line) b) Receiver Operator Characteristic curve, with False Positive Rate on x-axis and True Positive Rate on y-axis c) Model prediction output with segmentation masks.

### The model focuses on features inside the pathologies for classification

To understand what important features the model uses to predict the different types of plaques, we used Ablation-CAM^37^. The relationships that form between the layers of trained models such as our Mask R-CNN are non-linear and too complex to interpret directly; as a result, it is difficult to determine how the trained models make the classifications they learn to make. New tools such as Ablation-CAM have been developed to reveal to the human eye features in the image that the network learned to use to perform classifications. To implement Ablation-CAM, we first employed a gradient-free localization approach to obtain the feature maps. Then we ablated the feature maps randomly, and visualized the quality and accuracy of pixel-wise predictions (Fig. 6). After repeating these steps on all the feature maps, we obtained a heatmap of pixel areas that the Mask R-CNN learned to use to make its predictions. We found that the pixels that the Mask R-CNN learned to use to correctly classify each pathology were primarily within the pathology object itself. This finding indicates that the model bases its classifications on the objects themselves, rather than on surrounding features that could just be background noise. Ablation-CAM does not reveal exactly how the model makes its prediction, just what features/pixels of the object are most informative. We assume that the model uses variations in intensity and spatial distribution within the pathologies as reliable indicators of class, like a trained pathologist does.

**Figure 6.**
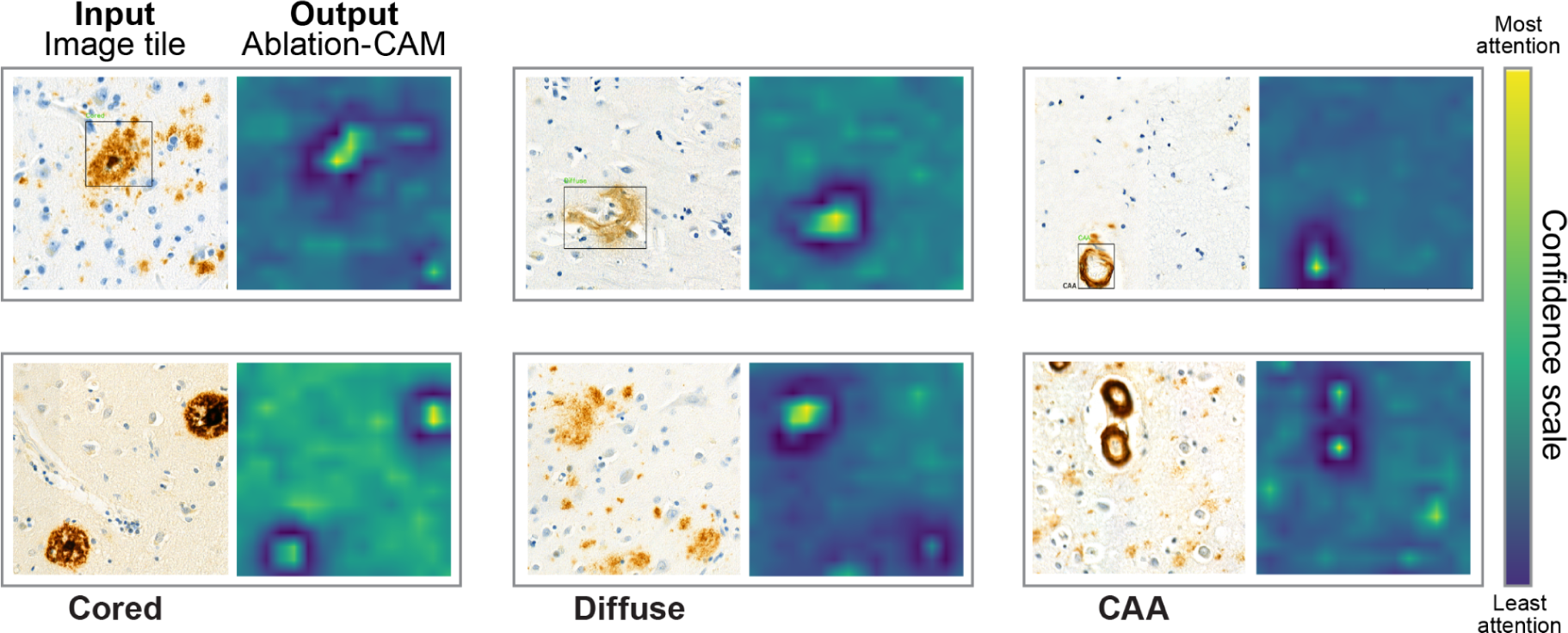
Ablation CAM output. Analysis of the Mask R-CNN trained to identify and classify different classes of amyloid pathology with the Ablation-CAM tool to interpret AI reveals image features that the network learned to drive the classification.

### Plaque segmentation map at Whole Slide Level

The determination of many types of neuropathology requires evaluation of labeled tissue samples at high spatial resolution, achieved with microscopy performed at high magnification. But context is also critical, and the clinicopathological significance of pathology may be apparent at multiple scales. For example, the major Braak staging system for AD is based on the extent of neurofibrillary tangle (NFT) pathology across different brain regions, rather than on microscopic features of the NFTs in a specific brain region^38, 39^.

To visualize the localization and distribution of various types of Aβ pathologies relative to macroscopic context features such as white matter (WM) and gray matter (GM), we used the positional information of each tile (1024 x 1024) from a WSI and stitched the tiles together to generate a segmented WSI heatmap (Fig. 7). To enhance visualization, segmented masks were overlaid on the individual plaques. Overlapping tiles were processed using the Non-Maximal Suppression algorithm^40^. To speed up the process, the stitching of tiles was done by replacing the original tiles in the memory location by the predicted tiles. This process took 7 minutes, 3 times less time than the initial break-down of WSIs into tiles.

**Figure 7.**
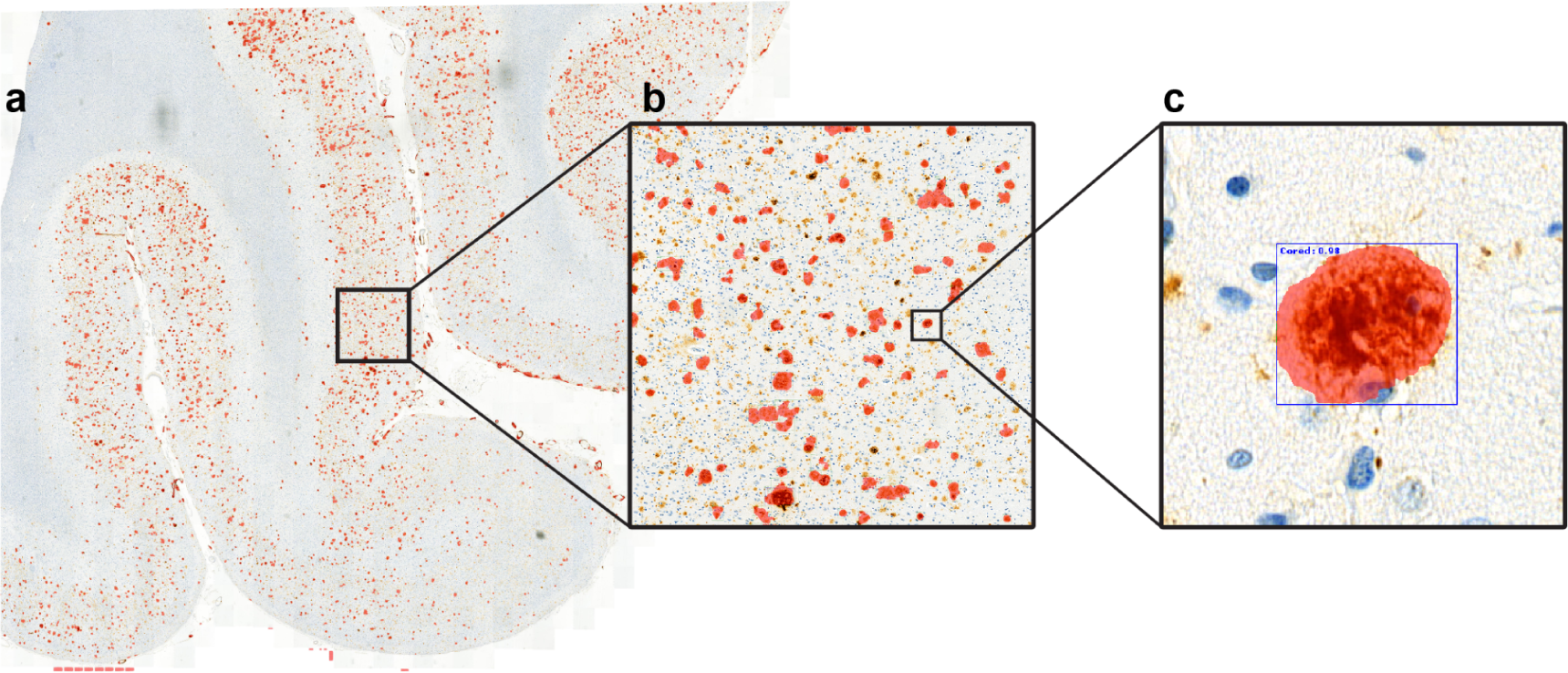
WSI plaque segmentation map a) Shows a segmented map of amyloid pathology at the WSI level. b) Zoomed in section of a WSI depicting the model predictions c) Shows the model segmented output for an individual core plaque. Red color mask is overlaid on the plaques to create this visualization.

The stitched-up map reveals that the majority of Abeta pathology is concentrated in the gray matter, aligning with the expected observations from high-resolution visual examination of stained slides by expert pathologists. This ability to rapidly stitch together the tiles along with their corresponding predictions allows for the visualization of macroscopic patterns in the distribution of pathology that were previously challenging or impossible to discern.

### Predictions from Mask R-CNN overlap with the CERAD ratings

The Consortium to Establish a Registry for Alzheimer’s disease (CERAD) has established standardized procedures for the evaluation and diagnosis of patients with AD^41^, including some of the best-validated tools to relate amyloid pathology to disease severity^24^. The CERAD ratings are calculated by evaluating the density of neuritic plaques in specific brain regions and categorizing them into sparse, moderate and frequent. We wondered if the automated measures of amyloid pathology made by the Mask R-CNN from postmortem patient brain samples correlated with the CERAD ratings of the severity of AD for those patients during life. To investigate, we applied our trained model to identify and quantify plaques in a test dataset comprising 86 Whole Slide Images (WSIs). The plaque quantification was compared with the CERAD ratings calculated by a trained pathologist, in the Middle Frontal Gyrus region of the brain. According to CERAD, the median value of neuritic plaques is 208 for “frequent”, 34 for “moderate”, and 10 for “sparse” rating. Our model predicted plaque counts that perfectly aligned with these ratings. To validate the correlation, we conducted a one-way ANOVA test, which revealed a significant difference between the means of the “moderate” and “frequent” groups (p < 0.0058).

We also found a positive linear correlation between the total number of neuritic plaques with cores predicted by the model and the average density of neuritic plaques assessed by the expert (Fig. 9). Furthermore, a Spearman Correlation performed between expert-based average neuritic density and model-predicted total neuritic plaques with cores yielded a value of 0.456, confirming a positive correlation between the two.

**Figure 8.**
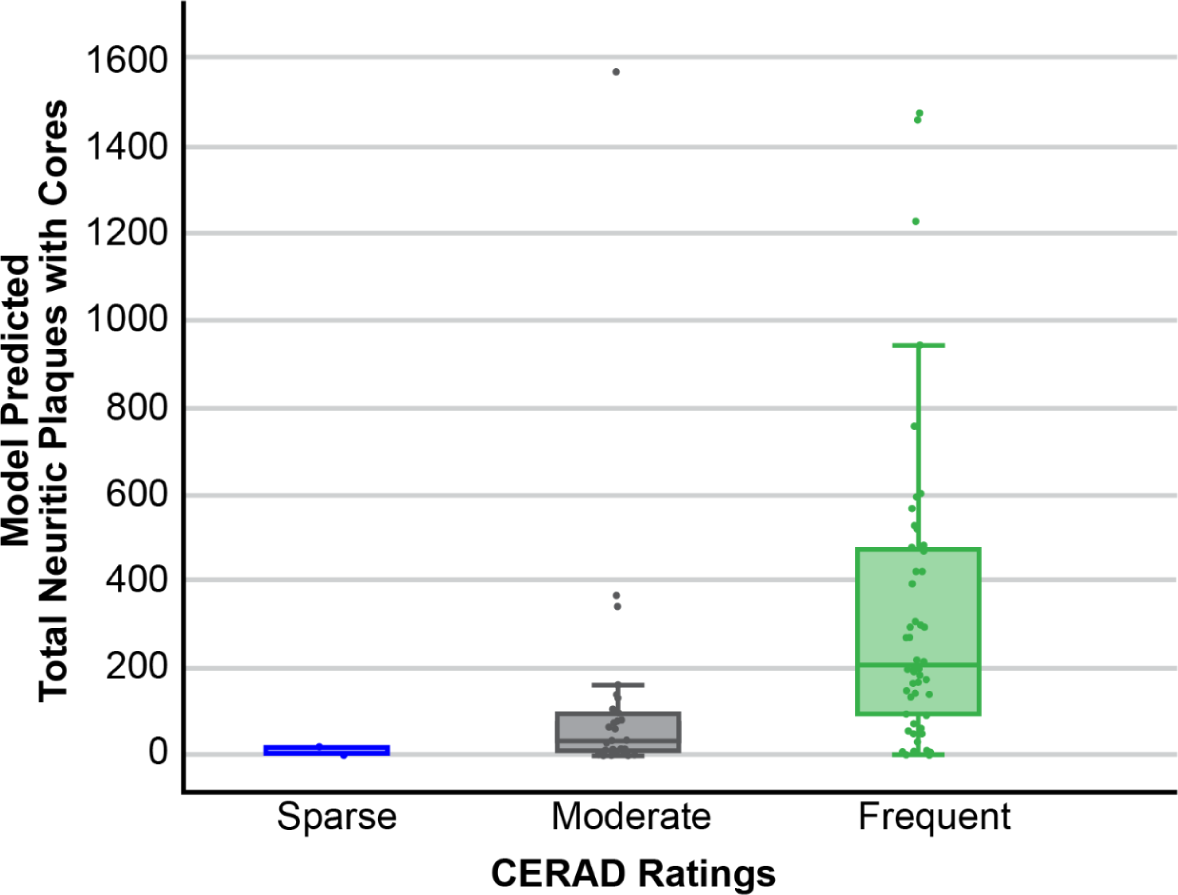
Quantification of amyloid pathology with the Mask R-CNN models correlates positively and significantly with the CERAD ratings of AD plaques for each patient. Box plot showing correlation of model predicted total neuritic plaques with cores, and the expert based CERAD ratings: Sparse, Moderate, and Frequent.

**Figure 9.**
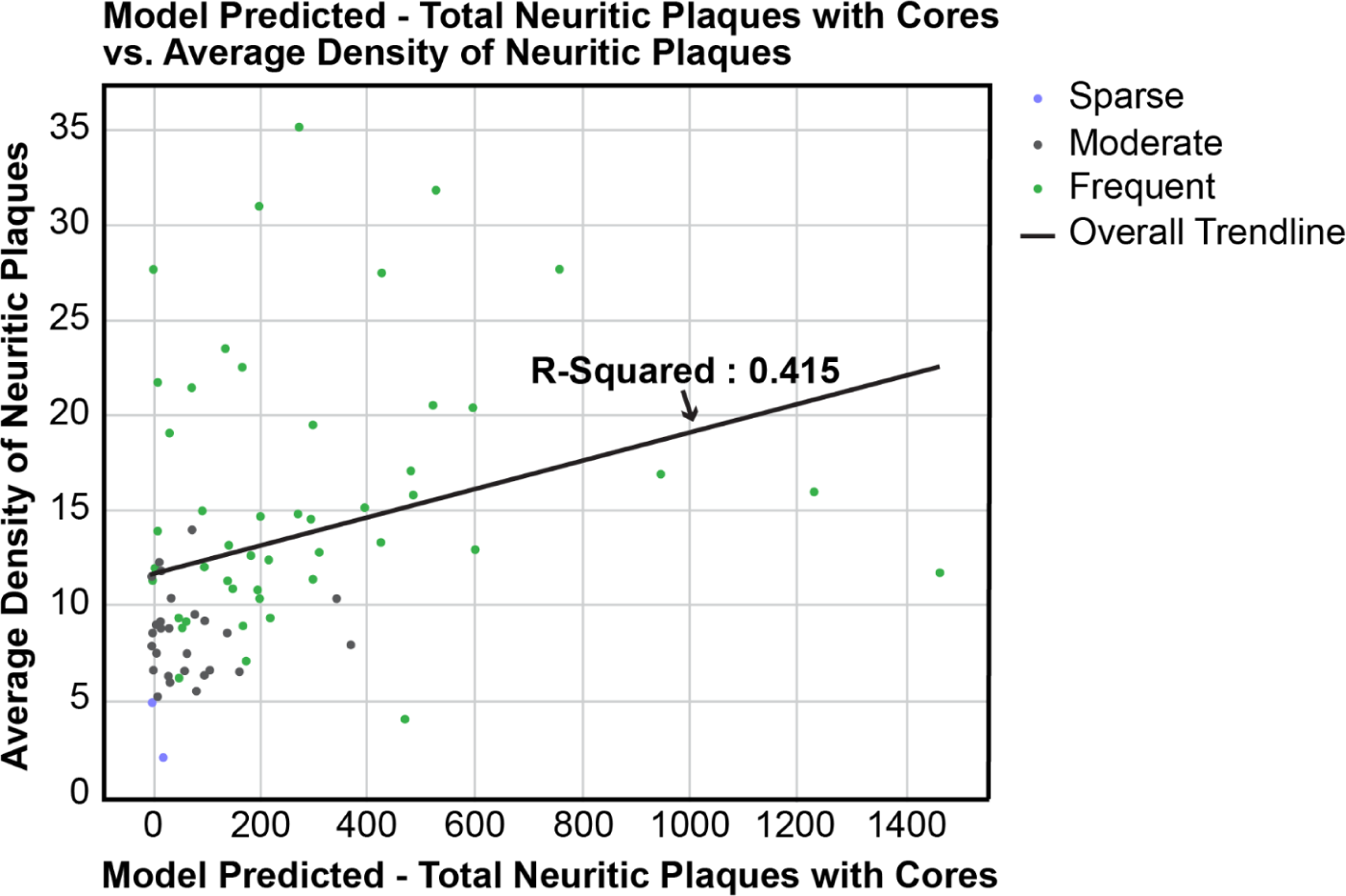
Quantification of amyloid pathology with the Mask R-CNN models correlates positively with the average density of neuritic plaques for each patient. Scatter plot showing model predicted neuritic plaques with cores on x-axis, and average density of neuritic plaques on y-axis. A regression line is fit having an R-squared value of 0.415

## Methods

### Ethics approval

These studies utilized tissue obtained from postmortem human brain specimens. All the specimens and WSIs are deindentifed.

### Case cohort and Sample preparation

Our dataset includes a total of 103 cases obtained from Mount Sinai Brain Bank, all of which are diagnosed with Alzheimer’s disease. A medical record review was performed on all cases using a structured instrument that records and codes all medical and neuropsychiatric conditions (based on ICD-9/10 and DSM IV/V codes), all medications (including doses and duration), the severity of dementia (based on clinical dementia rating), neuropsychiatric symptoms (NPS), family history, available laboratory and neuroimaging study results and sociodemographic variables such as level of education and languages. Brain sections from middle frontal gyrus were prepared from each AD case in paraffin-embedded blocks and immunolabeled with anti-β amyloid (4G8). Neuropathological diagnostic criteria for AD are based on CERAD, which includes quantification of neuritic plaques (NP) in the middle frontal gyrus brain region.

### Dataset preparation

The dataset consists of 4210 tiles of size 1024 x 1024 pixels obtained from 17 WSIs and containing a classification label and a pixel-level segmentation mask annotated by expert pathologist in QuPath Software^31^. To provide more variation in the training cycle, objects in the dataset were not centered within the tiles and the dataset was augmented using various techniques shown in Table 3 from the Albumentation library^42^. The dataset was divided into a training dataset consisting of 3188 tiles extracted from 10 whole slide images (WSIs), and a validation dataset containing 1022 tiles obtained from 7 WSIs.

**Table 3.** Data augmentation techniques from Albumentation.

|  |
| --- |
| Vertical Flip (p = 0.5) |
| HorizontalFlip (p = 0.5) |
| Blur(blur_limit = 3) |
| OpticalDistortion() |
| HueSaturationValue() |
| RandomRotate90() |

### Pipeline

The WSI is loaded into QuPath at 20X magnification. The resolution is set to 1 micron per pixel (MPP) and a grid is added to the Image View, where each grid is 1024 x 1024 pixels. Annotation is performed on grids that have Cored plaques, Diffuse plaques and CAA by a set of expert neuropathologists. Pixel-level segmentation of the plaque regions is done with a magic wand tool. During annotation a confidence score is added, to check for annotation interrater variability.

The confidence score values range from 1-3, where 1 means possible, 2 probable and 3 definite. The annotations are written into a custom JSON format containing all the important polygon regions and their corresponding labels. In the training phase the model is fed only the tiles annotated by experts containing objects of interest.

In the testing phase, the model is fed only the part of the WSI that contains tissue, which we determine by downscaling the WSI by a factor of 4 and then utilizing Otsu thresholding to generate a contour mask as shown in Figure 10.

**Figure 10.**
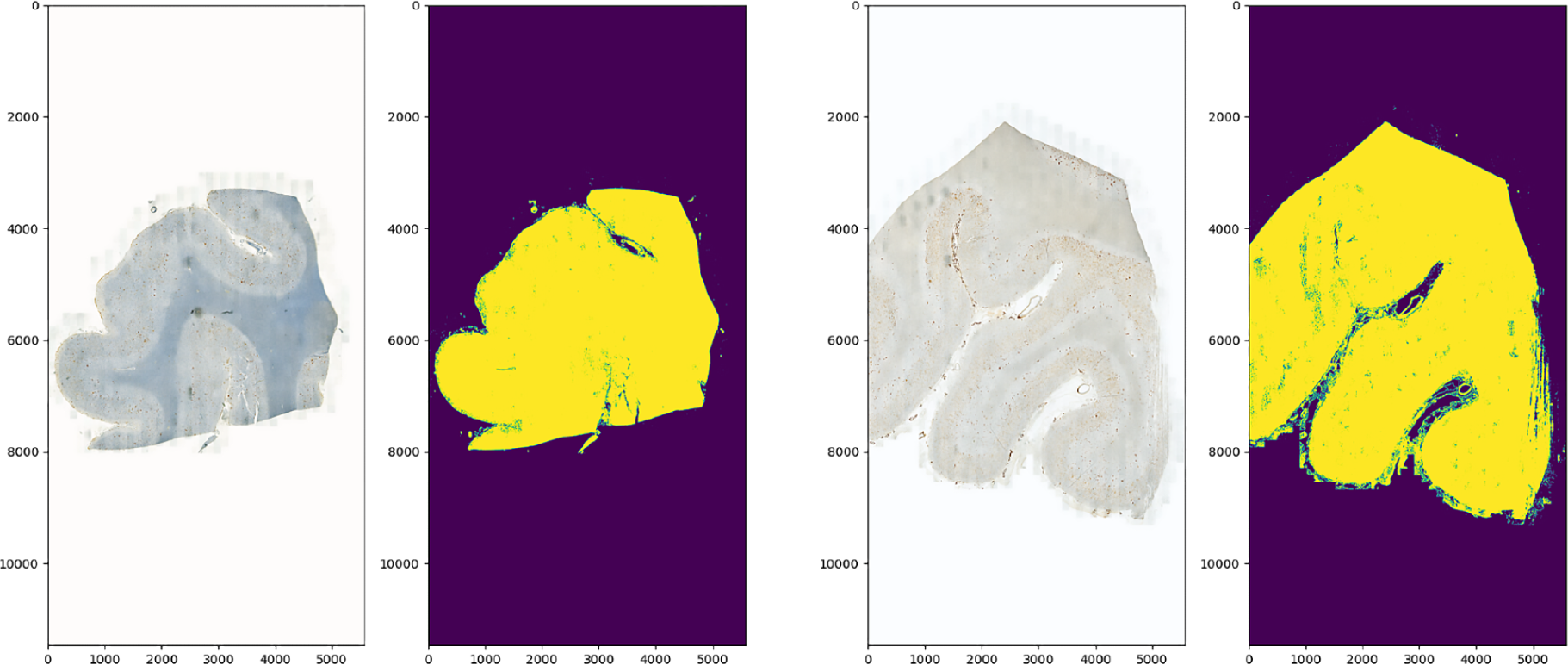
WSI tissue segmentation using Otsu thresholding.

### Model Development and Training

Pytorch framework is used for the development of the model. All the models are trained on a single NVIDIA Quadro RTX 5000 GPU. The architecture is based on the Mask Regional Convolutional Neural Network (Mask R-CNN)^30^. The Mask R-CNN model is split up into three stages. The first stage uses a Feature Pyramid Network (FPN) to extract the features from an input image at different scales (Figure 11.1). A 1 X 1 convolution filter is applied to all the feature maps from the corresponding image scales and the resulting features maps are averaged to form the final feature map. This step ensures that the feature map contains important information from both low resolution and high resolution images for a given object of interest. We use the pre-trained weights from ResNet50 as our backbone architecture for the FPN model.

The second stage (Fig. 11.2) includes a lightweight neural network called the Region Proposal Network (RPN) that scans the feature map generated by an FPN to propose regions that may contain a plaque. RPN relies on anchor points and anchor boxes. Every point in a feature map is an anchor point. We generate anchor boxes for every anchor point. The anchor boxes help us to map the features from the feature map to the raw image location in the input image. A set of 9 anchor boxes are generated for each anchor point based on different combinations of scale and aspect ratios.

**Figure 11.**
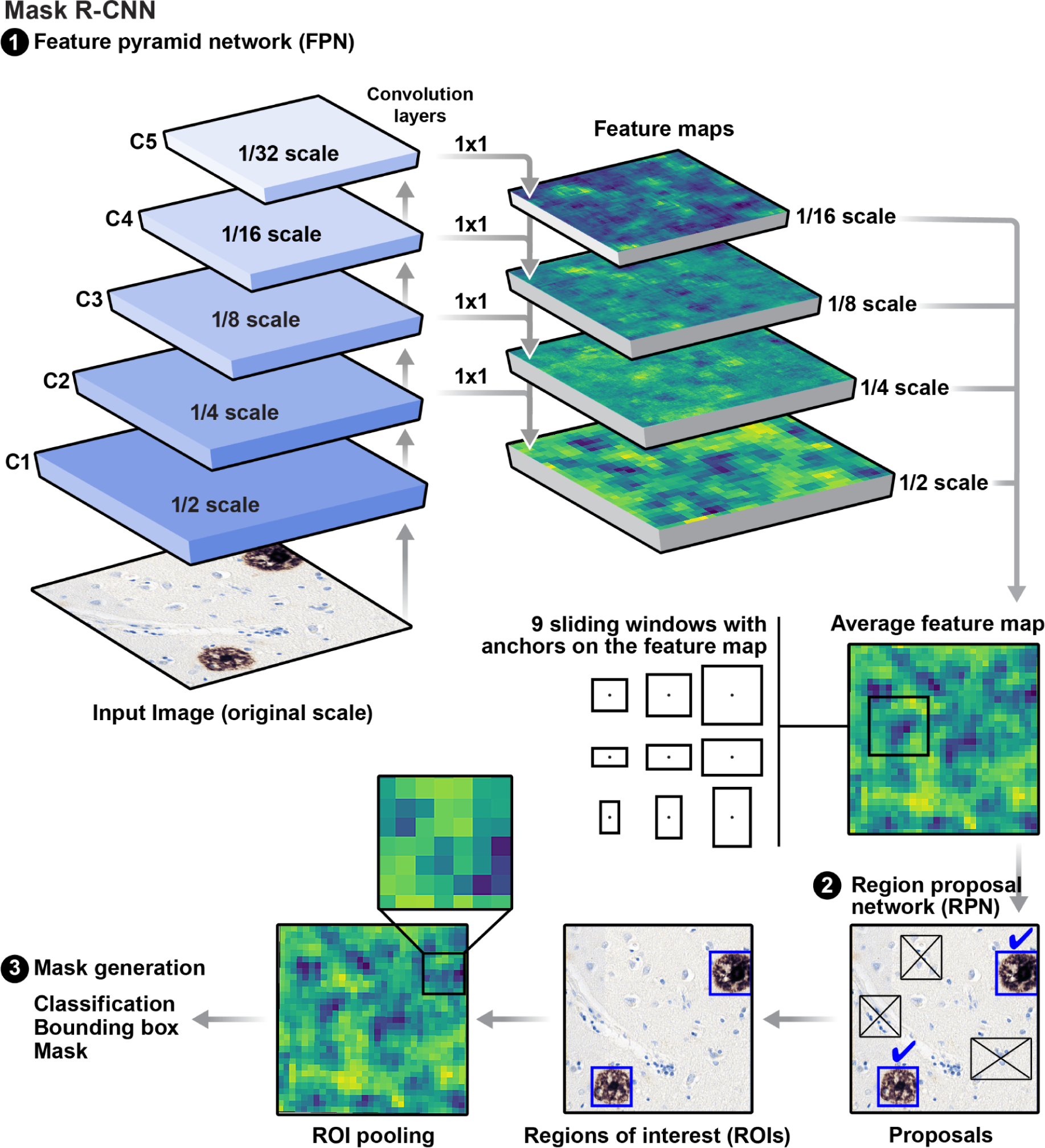
Model architecture based on Mask R-CNN Feature Pyramid Network with ResNet50 as the backbone architecture. Feature maps are generated from images at different scales and are used for downstream prediction.

To get good quality anchors, a Non Maximal suppression algorithm is used where all the anchors whose intersection of union (IoU) with the ground truth (GT) box is below 0.7 are discarded. The RPN takes these anchors and uses a fully connected layer to give two outputs: (1) - binary class - whether an anchor contains an object or not and (2) the bounding box regressor representing the delta in x, y, w, and h that are required to come closer to the GT boxes.

In the third stage (FIg. 11.3), the network takes in the regions proposed by RPN as input and uses Region of Interest (ROI) align pooling to extract the corresponding feature map without quantization. This feature map is sent to a set of CNN’s for generating pixel-wise classification and segmentation masks. We use the following loss function

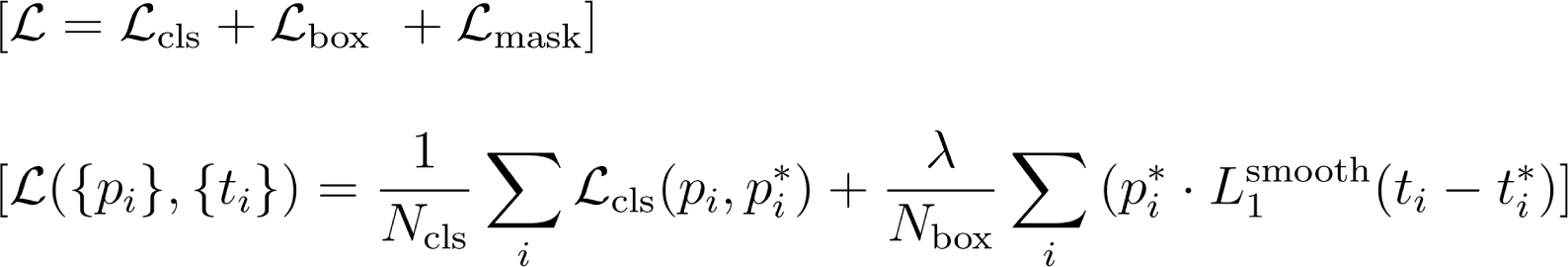

where 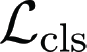 is the log loss function over two classes. For the box loss, we can easily translate a multi-class classification into a binary classification by predicting a sample being a target object versus not as 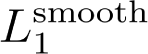 loss.

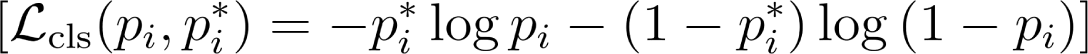

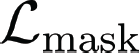 is defined as the average binary cross-entropy loss, only including *k*th mask if the region is associated with the ground truth class *k*.

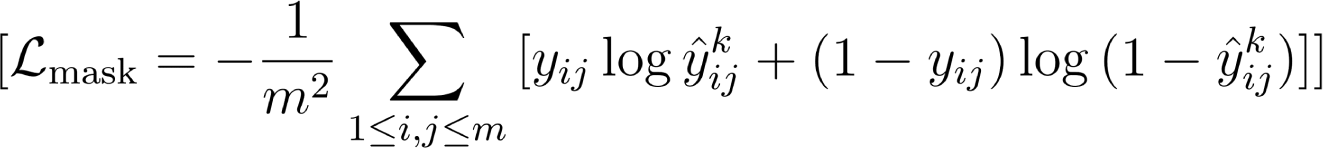

Where *y_ij_* is the label of a cell (*i, j*) in the true mask for the region of size $m\times m$; 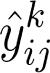 is the predicted value of the same cell in the mask learned for the ground-truth class *k*.

The model is trained for 100 epochs with a batch size of 10. The evaluation is done every 10 epochs. We use Stochastic Gradient Descent as the optimizer of the model with a learning rate of 10^-3^.

### Implementation of Ablation CAM

To understand further what important features the model uses to predict the different types of plaques, we use Ablation-CAM^37^. In this technique, we employ a gradient-free localization approach to visualize the predictions from the model (Fig. 12). The feature maps generated from the FPN model are randomly ablated to check how they affect the pixel-wise predictions. y^c^ is the probability of class C. y^ck^ is the probability of class C after ablation. If a certain pixel is important, then the difference between y^c^ and y^ck^ will be large. This is done repeatedly on all the feature maps. The final output of Ablation-CAM is a heatmap of pixel areas that are critically important for the prediction.

**Figure 12.**
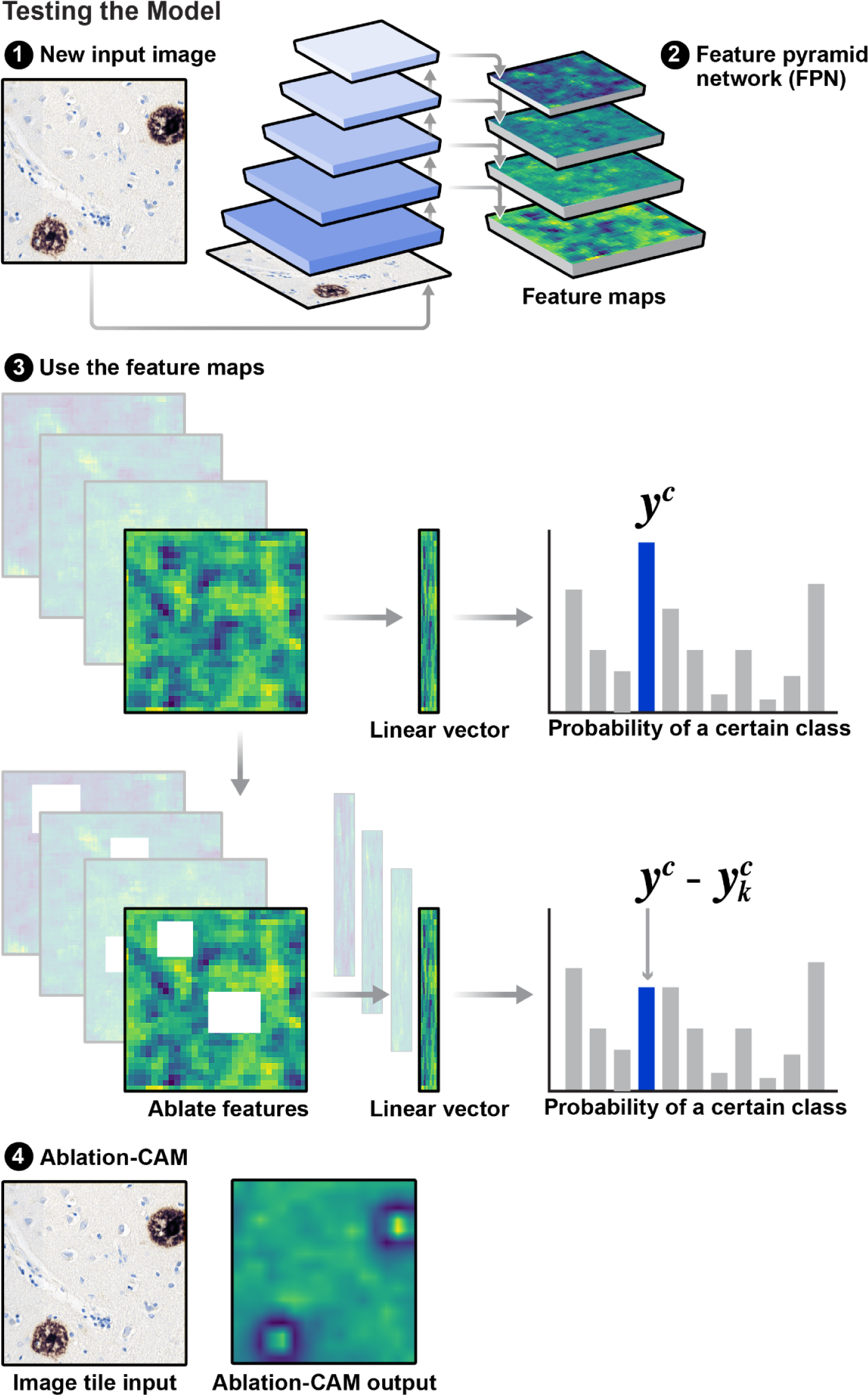
Model Explainability using Ablation CAM.

## Discussion

In this study, we developed a scalable computational pipeline to batch-process large numbers of WSIs in high throughput and facilitate large-scale, unbiased, comprehensive and quantitative assessment of neuropathology. We then used the pipeline to train classifier models that could identify and classify with human-level accuracy three types of amyloid pathology: cored plaque pathology, diffuse plaque pathology, and amyloid angiopathy. In addition to assessing the quality of the models with standard metrics, we tested the models on a dataset from an independent brain bank not used during model training. The prediction accuracies of the models indicate that they learned generalizable features of these three classes of amyloid pathology. With these models, we showed that automated quantification of amyloid pathology in a postmortem patient material was significantly correlated with the severity of AD during that patient’s life as measured by CERAD score.

Several advances in the design of the computational pipeline contributed to its successful application. The decision to leverage the GPU architecture and use 1024 x 1024 crops instead of the more widely used 256 x 256 crops improved performance two ways. First, based on feedback from our expert neuropathologists, the larger crops provided critically needed additional context to properly annotate images for the training dataset. The large crops also led to a ∼5-fold increase in throughput, reducing the processing time for a WSI from 109 minutes to 21 minutes.

Another advance was the use of a Mask R-CNN architecture. The incorporation of the RPN sped up the throughput of the pipeline by narrowing the regions to analyze fully to those with a high probability of containing neuropathology features of interest. The Mask R-CNN also is capable of performing simultaneous identification and segmentation of neuropathological objects, which enables the investigator to go beyond simple counts of pathology and perform deeper analysis of the objects including size, shape, texture etc. While we did not use this asset of Mask R-CNN in this study, we anticipate it could ultimately reveal sub-features of pathology that correlate with specific features of the patient.

We showed that the Mask R-CNN architecture could be used to train models to accurately classify cored plaque, neuritic plaque and cerebral amyloid angiopathy pathology with human level performance. Precision-recall analysis showed that these models performed well by standard measures, and k-fold cross validation accuracy indicated the models performed stably over different segments of data, suggesting the models were not overfit. With the interpretable AI tool, Ablation-CAM, we showed that the Mask R-CNN models had learned to make accurate predictions of the presence and class of amyloid pathology based on characteristic patterns of pixels within the pathological objects, adding confidence that the Mask R-CNN had learned essential features of distinct amyloid pathologies.

The ability of the Mask R-CNN to use amyloid β labeling to correctly identify and distinguish classes of hallmark neuropathology and yet ignore unspecific amyloid β labeling highlights the tremendous power of machine learning. In simple terms, with enough ground truth examples of objects that the human annotator determines the DL networks should attend to as well as ignore, the training process iteratively remodels the weights assigned to image features until it can make accurate predictions. In contrast to simple image analysis approaches that rely on defining foreground pixels by setting a threshold based on their intensity and defining objects based on groups of foreground pixels that are certain size or area, DL networks are capable of recognizing much more complex spatial patterns and at multiple scales, formed by variations in pixel intensity and distribution. This ability accounts for the dramatic power of DL to resolve sometimes subtle differences between objects and to achieve human level accuracy. In certain ways, the DL network is able to more fully exploit the data in images than humans eyes can since it can easily take full advantage of the full range of pixel intensity values within an image, which can range from 16,383 to 65,546 distinct gray values for WSIs digitized with 14-bit or 16-bit cameras. That contrasts with only 256 gray values available to humans observing images on standard computer monitors. The fact that DL networks can resolve patterns down to the pixel level may also explain why DL networks can make accurate predictions based on digitized images collected at 20x magnification and do not require the greater spatial resolution afforded by images collected at 40x magnification. The deep layer structure enables the network to extract simple image features in the initial layers of the architecture and then iteratively learn to find and weight more complex patterns formed from those simple features that improve prediction accuracy. Unsurprisingly, Ablation-CAM revealed that the DL network had learned to make accurate classifications using a set of simple and complex image features similar to those used by human annotators to create the ground truth dataset.

Importantly, the models were tested on neuropathology data from an independent cohort of Alzheimer’s disease patients prepared by the University of California, Davis Brain Bank. Few published models have been tested beyond the dataset on which they were trained, in part because it has been difficult to obtain publicly available annotated AD neuropathology datasets. Unfortunately, without testing algorithms on independent datasets, it is difficult to judge their generalizability. This is a concern with digital pathology because of the many potential ways in which isolated training datasets may have unique features that could be learned by DL networks and would limit their generalizability. For example, different brain banks often have slightly different sample preparation techniques, and the intensity of labeling can vary from sample to sample, even within the same brain bank. Other studies have shown that the assessments made by expert pathologists also can vary^43–47^. To the extent that this is true, algorithms could learn from annotations to accurately mimic the predictions of one pathologist but their accuracy would fall if they were applied to a different dataset annotated by a pathologist with different judgments. Since we tested the generalizability of our algorithms on a dataset generated by an independent brain bank on an independent cohort of patients, all the potential issues described above could be relevant. In addition, the independent dataset came in the conventional form of 256 x 256 image crops. To be able to assess the performance of our algorithms on those data, we first needed to resize the crops to 1024 x 1024, an operation that almost certainly caused some loss of detail and sharpness in the resultant images. Yet our models accurately predicted the three types of amyloid pathology in these resized images. Unsurprisingly, the performance of the models was not as high on this new dataset as it was on the validation dataset. Nevertheless, the fact that our models were performative strongly indicates that they had learned generalizable features of amyloid pathology that span different AD patients, brain banks and neuropathologists.

The trained models were then used to determine whether and to what extent the quantification of amyloid pathology in postmortem material from patients correlates with the severity of AD during life, as measured by CERAD scores and we found there was a positive and significant correlation between the expert estimated plaque burden and the unbiased automated quantitative measures of amyloid pathology. An important future direction will be to leverage the capability of the Mask R-CNN to segment pathological objects to determine whether more specific features of certain types of amyloid pathology have an even stronger predictive value for AD severity or specific symptoms of the AD patients.

In sum, this work demonstrates the power of the pipeline to analyze large sets of WSIs, which will enable the exploration of correlations between neuropathological features of AD and an array of other patient data including patient-specific clinical features and genotypes. In turn, the ability to perform such large-scale explorations could help reveal how genetic variants drive the development of different types of CNS pathology through cell-specific and autonomous effects as well as through interactions between different cell types and between cells and hallmark pathology.

## Acknowledgements

We wish to express our sincere gratitude to Dr. Brittany Dugger who generously made available digitized images of neuropathology of AD patients from the University of California, Davis Brain Bank for testing the generalizability of our algorithms. We would like to thank Dr. Laura Parkkinen for providing additional neuropathology input and expertise. We also wish to thank Françoise Chanut for editorial support and Kelley Nelson and Gayane Abramova for administrative support with manuscript preparation. This work was made possible primarily with support by the National Institutes on Aging (R01 AG067025) and the National Library of Medicine (R01 LM013617). Additional support was from the National Institutes of Neurological Disorders and Stroke (R01 NS124848) and the Koret Foundation Biomedical Research Program in Artificial Intelligence.

